# Single–gene knockout of RNLS or HIVEP2 are insufficient to protect β–cell spheroids from allo– and xeno–rejection

**DOI:** 10.1101/2025.07.30.667633

**Authors:** Ismail Can Karaoglu, Arda Odabas, Tamer Onder, Seda Kizilel

## Abstract

Type 1 diabetes can be cured by β–cell replacement in principle, yet recurrent autoimmunity and transplantation barriers rapidly destroy implanted cells. Genome–wide CRISPR screening by Cai *et al*. highlighted RNLS and HIVEP2 as candidate genes, but their value outside an autoimmune setting is unknown. Here, it was evaluated whether single-gene knockout of RNLS or HIVEP2 could similarly protect β-cell grafts against allo- and xenogeneic rejection. Murine β–TC–6 and human EndoC–βH1 cell lines were genetically edited using CRISPR-Cas9 to knockout RNLS or HIVEP2, and editing efficiencies were confirmed via T7 endonuclease I assay and TIDE analysis. Functional characterization indicated that RNLS deletion modestly impaired glucose-stimulated insulin secretion in murine cells, whereas HIVEP2 deletion showed no functional alterations in either cell line. For *in vivo* assessment, genetically edited β-cell spheroids were subcutaneously transplanted into CD-1 mice to model allo- (murine β-cells) and xenogeneic (human β-cells) rejection scenarios. Bioluminescence imaging revealed no protective effects of RNLS or HIVEP2 deletion, with grafts from both knockout groups displaying identical rejection kinetics compared to controls. These findings indicate that single-gene deletions of RNLS or HIVEP2 are insufficient for conferring meaningful protection against allo- or xenorejection, highlighting the necessity for combinatorial genome editing strategies or complementary biomaterial-based immunomodulation to achieve effective and sustained β-cell graft survival.

## Introduction

Type 1 diabetes (T1D) arises from autoimmune killing of the insulin–producing β–cells of the pancreatic islets, forcing patients into lifelong exogenous–insulin therapy [1–4]. This exposes them to hypoglycemia and chronic complications such as heart disease, stroke, nerve damage, and kidney failure [5–9]. β–cell replacement therapy offers curative potential, yet transplanted grafts rejected rapidly to both recurrent autoimmunity and the de novo allogeneic or xenogeneic immune responses of the recipient. Current clinical protocols rely on broad immunosuppression, which incompletely protects grafts and carries substantial infectious and oncogenic risk [6, 10–18]. Consequently, engineering β–cells that evade immune attack while preserving host immunosurveillance is a priority in the field [19–25].

Gene–editing has emerged as a powerful route to such “stealth” cells. Ablation of classical HLA class I and II molecules or over–expression of CD47 demonstrably reduces T–cell recognition, but complete removal of HLA can provoke natural–killer (NK)–cell activation and compromise pathogen defense [26–30]. A complementary strategy is to identify β–cell–intrinsic genes whose loss confers resistance to immune injury without disabling antigen presentation. In this context, a genome–wide CRISPR screen performed by Cai et al. in non–obese–diabetic (NOD) β–cells uncovered two such loci, RNLS and HIVEP2, whose disruption enabled mutant cells to survive autoimmune assault *in vivo* [31]. RNLS encodes Renalase, a secreted flavoprotein oxidase that metabolizes atypical NAD(P)H isomers and has been linked to redox homeostasis and protection from cytokine–induced apoptosis. Its deletion in β–cells attenuate ER–stress signaling and dampens autoreactive CD8^+^–cell activation [32]. HIVEP2 (Schnurri–2) is a zinc–finger transcription factor that binds κB motifs, modulates NF–κB–driven transcription and restrains pro–inflammatory gene expression [33]. Although both knock–outs (KOs) provided an autoimmune survival advantage in the screen, their capacity to protect against the distinct mechanisms underlying allogeneic rejection (MHC mismatch and indirect antigen presentation) or xenogeneic rejection (innate immunity, complement and xeno–reactive antibodies) has never been tested.

This study aimed to test the hypothesis that single-gene knockout strategies targeting immune-regulatory and cellular protection genes, specifically, RNLS or HIVEP2, would improve β-cell survival by conferring resistance against allo- and xenogeneic immune rejection. These genes were selected based on their previously identified roles in regulating immune-mediated cellular stress responses and apoptosis pathways. Specifically, the hypothesis here is that knocking out RNLS would protect β-cells by reducing endoplasmic reticulum stress-induced apoptosis, whereas knocking out HIVEP2 might enhance β-cell resistance by altering immune recognition and cytokine responses. RNLS was identified as a promising target gene in genome-wide CRISPR screening, linked primarily with apoptosis regulation under inflammatory stress conditions frequently observed in autoimmune diabetes. It was hypothesized that knocking out RNLS would mitigate stress-induced β-cell apoptosis, thus enhancing graft survival. Similarly, HIVEP2 was chosen due to its known regulatory roles in immune responses, which could protect against autoimmune attacks by modulating cytokine signaling and immune-cell interactions.

Here, it was investigated whether deleting RNLS or HIVEP2 prolongs β–cell survival beyond the autoimmune context. Murine β–TC–6 and human EndoC–βH1 cells were first edited with pLentiCRISPRv2 constructs targeting RNLS or HIVEP2, and indel formation was verified by T7 endonuclease assay and TIDE analysis. Then each line was aggregated into uniformly sized (~150 µm) spheroids on agarose–coated ultra–low–attachment dishes. Functional testing revealed that RNLS KO modestly reduced glucose–stimulated insulin secretion in mouse cells, whereas HIVEP2 KO was functionally neutral in both species. To assess graft fate in immunocompetent hosts, luciferase–labelled spheroids were embedded in Matrigel and implanted subcutaneously into CD–1 mice, modelling either allogeneic rejection (β–TC–6 → CD–1) or xenogeneic rejection (EndoC–βH1 → CD–1). Longitudinal bioluminescence imaging demonstrated indistinguishable clearance kinetics among RNLS–, HIVEP2– and non–targeting grafts in both settings. Our findings therefore show that single–gene disruption of RNLS or HIVEP2 is insufficient to protect β–cell grafts from allo– or xeno–immunity. These results set practical limits on the utility of these loci as stand–alone edits and underlining the need for combinatorial genome engineering or complementary immunomodulatory biomaterials to achieve clinically durable immune evasion.

## Results and Discussion

### β–TC–6 and EndoC–βH1 cell spheroid optimization

Prior to the gene–editing experiments, 3D β–cell culture was optimized to obtain size–controlled, high–viability spheroids that more closely mimic the cell–cell interactions of native islets. The first approach relied on the hanging–drop technique. Single–cell suspensions of β–TC–6 cells were adjusted to 300, 500, or 1000 cells per 30 µL droplet, arrayed on the inverted lids of Petri dishes and cultured for 72 h at 37 °C under 5 % CO_2_. During this period, cells go to the bottom of the droplet due to gravity, where they aggregated into spheroids. Bright–field microscopy revealed a direct relationship between initial seeding density and final aggregate size. Droplets seeded with 300 cells produced spheroids averaging 120 µm in diameter, whereas those initiated with 1000 cells routinely exceeded 200 µm. Live/Dead staining demonstrated that aggregates below 150 µm were uniformly viable, while the larger spheroids displayed a central necrotic zone, consistent with oxygen– and nutrient–diffusion limitations reported in the literature (Figure 1a and b) [34]. Although the method can be considered as simple, the hanging–drop format proved labor–intensive, and difficult to scale beyond a few hundred spheroids per lid, making it impractical for the cell quantities required for transplantation.

**Figure 1.**
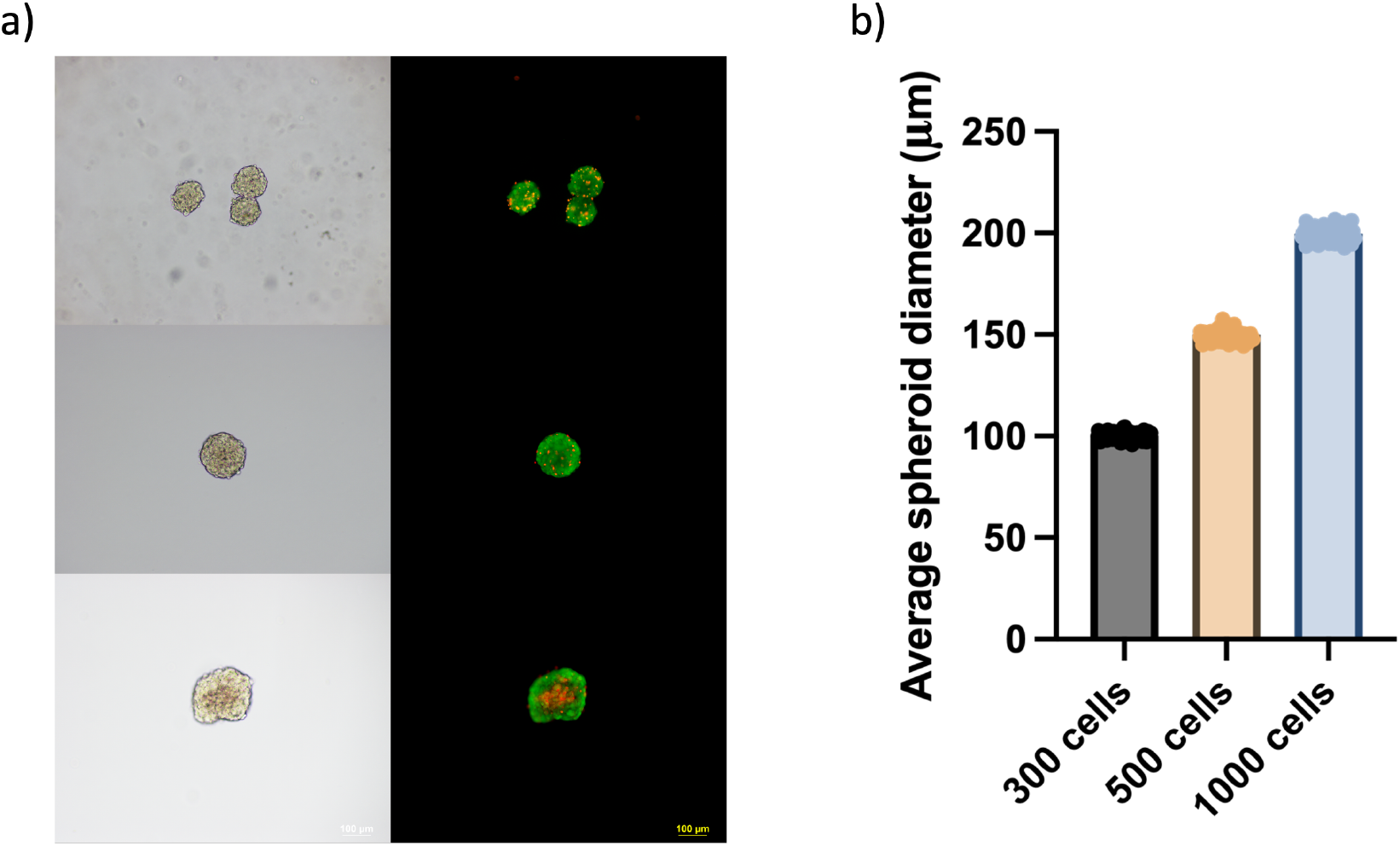
Hanging–drop optimization of β–TC–6 spheroid size and viability. **a)** Representative bright–field (left) and Live/Dead (right; fluorescein diacetate, green = viable; propidium iodide, red = non–viable) images of spheroids generated from 300, 500 and 1000 cells per 30 µL hanging drop (top to bottom). Spheroids seeded with 1 000 cells routinely exceed 200 µm and display a central necrotic core, whereas aggregates formed from ≤ 500 cells remain compact and uniformly viable. (Scale bar = 100 µm) **b)** Quantification of average spheroid diameter (mean ± SD, n = 100 independent droplet sets per condition). Diameter increases proportionally with initial cell number, emphasizing the need to limit seeding density to < 500 cells to maintain physiologically relevant (< 150 µm), necrosis–free aggregates.

To overcome these constraints, a suspension culture protocol was developed using home–made ultra–low–attachment dishes. Petri dishes were coated with 1 % (w/v) agarose solution and allowed to solidify, creating a non–adhesive surface that prevents cell attachment. 4 × 10^6^ single cells were added to each dish in 8 mL of complete medium and cultured for 72 h on an orbital shaker set to 50 rpm. Under these conditions, cells aggregated spontaneously into hundreds of uniformly round spheroids (Figure 2a and 2b). Image analysis of 100 aggregates showed a narrow size distribution centered at 140 ± 8 µm, while confocal Live/Dead staining confirmed ≥ 95 % viability throughout the spheroid cross–section (Figure 2c). Notably, the suspension approach yielded a higher number of spheroids per dish compared to the hanging–drop method, making it the preferred method for downstream experiments [Costa et al. 2018].

**Figure 2.**
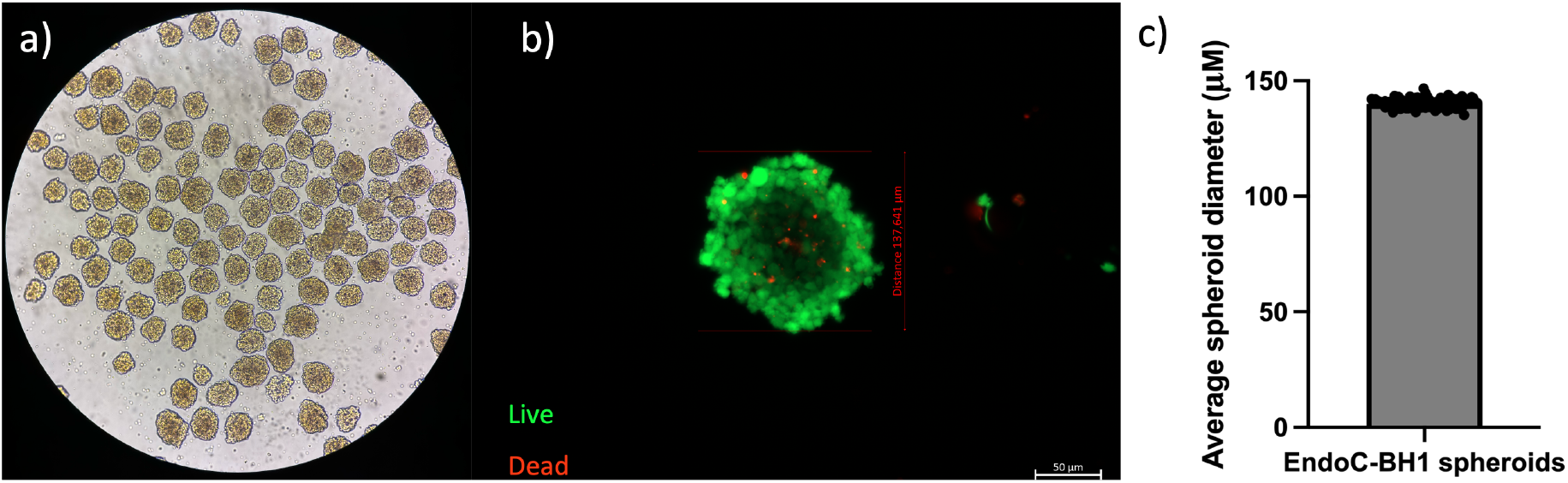
Suspension–culture generation of uniformly sized, viable EndoC–βH1 spheroids. **a)** Bright–field overview of the agarose–coated ultra–low–attachment dish after 72 h of orbital shaking (50 rpm) with 5 × 10^6^ human EndoC–βH1 cells, showing hundreds of compact aggregates. **b)** Live/Dead confocal image of a representative spheroid stained with fluorescein diacetate (green, viable) and propidium iodide (red, non–viable). The absence of red signal confirms minimal core necrosis. Scale bar = 50 µm. **c)** Mean spheroid diameter determined from 100 aggregates (140 ± 8 µm; mean ± SD), illustrating the narrow size distribution achieved with the shaking suspension method.

Given its superior throughput, precise size control, and absence of a necrotic core, the agarose–based suspension culture was selected for all subsequent CRISPR/Cas9 editing, functional assays, and transplantation studies. While the hanging–drop data provided a valuable reference for correlating spheroid diameter with diffusion limits, this information guided our decision to maintain aggregate diameters below 150 µm in all subsequent experiments.

### T7 endonuclease I (T7E1) and Tracking of Indels by Decomposition (TIDE) analysis

To perform targeted gene knockouts, three guide RNAs (gRNA) for RNLS and HIVEP2 genes were selected from the Brie and Brunello databases, including the guide RNA previously validated by Cai *et al*. as a control for RNLS knockout [31]. As an internal control for functional knockout validation, the insulin gene (Ins2 for mouse, INS for human) was targeted in both cell lines. Each selected guide RNA was cloned into the pLentiCRISPRv2_puro plasmid, and successful insertion was confirmed by Sanger sequencing. Lentiviral particles were then produced by co–transfection of HEK293T cells with the guide RNA–cloned pLentiCRISPRv2_puro plasmids along with packaging (psPAX2) and envelope (VSV–g) plasmids. GFP–encoding lentivirus was produced similarly and used as a control to monitor infection efficiency. Following lentiviral production, β–TC–6 and EndoC–βH1 cell lines were infected with lentiviruses encoding gRNAs targeting RNLS, HIVEP2, and INS. After antibiotic selection to enrich for successfully transduced cells, cell pellets were collected and used for subsequent validation assays, including quantitative PCR (qPCR), genomic DNA isolation for T7 endonuclease I (T7E1), Tracking of Indels by Decomposition (TIDE) assays to confirm successful genomic editing at the intended target sites.

To validate successful genome editing in β–TC–6 cells, T7E1 mismatch assays were conducted for the Hivep2, Ins2, and Rnls gene targets. Distinct cleavage products were observed for gRNA2 targeting HIVEP2, all three gRNAs targeting INS2, and gRNA3 targeting RNLS, confirming the presence of CRISPR/Cas9–induced insertions and deletions (indels). No cleavage bands were detected in the corresponding non–targeting (NT) control lanes, confirming editing specificity (Figure 3a). Consistent with the T7E1 results, TIDE analysis of HIVEP2–edited β–TC–6 cells using gRNA2 revealed editing efficiency of approximately 43.7%, further validating robust gene disruption at the target site (Figure 3b).

**Figure 3.**
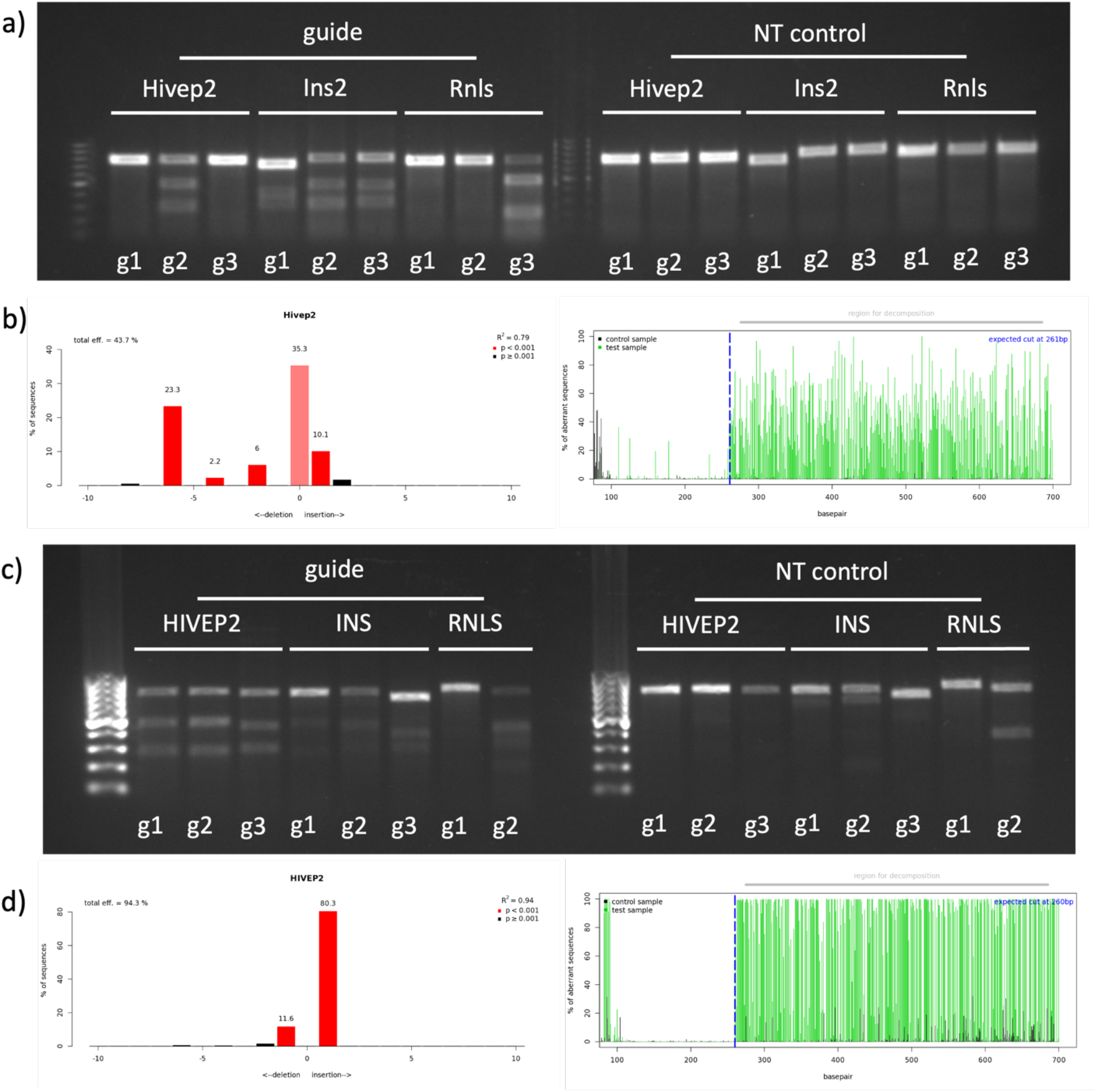
Genomic validation of CRISPR/Cas9 editing in mouse and human β–cell lines. **a)** T7E1 assay of genomic PCR products amplified from β–TC–6 cells transduced with three gRNAs targeting Hivep2, Ins2 or Rnls (left), alongside the corresponding non–targeting (NT) controls (right). Cleavage products visible for Hivep2–g2, all three Ins2 guides and Rnls–g3 confirm formation of indels, whereas NT lanes show only full–length amplicons. **b)** Tracking of Indels by Decomposition (TIDE) analysis of the most efficient mouse guides. Left: indel frequencies. Right: representative TIDE decomposition. **c)** T7E1 analysis of EndoC–βH1 cells edited with guides against HIVEP2, INS or RNLS. Pronounced cleavage is evident for all three HIVEP2 and INS guides, and modestly for RNLS–g2, whereas NT controls are uncleaved. **d)** TIDE quantification in human cells. Left: indel percentage. Right: representative TIDE decomposition.

In parallel, genome editing in EndoC–βH1 cells was assessed using the same approach. T7E1 assays revealed clear cleavage products for all three gRNAs targeting HIVEP2, INS, and gRNA2 targeting RNLS, indicating efficient CRISPR–mediated gene knockout events (Figure 3c). TIDE analysis of the HIVEP2 locus edited with gRNA1 demonstrated a high indel frequency of approximately 94.3%, corroborating the T7E1 data and confirming moderate editing efficiency in human β–cells (Figure 3d).

### Insulin production and glucose stimulated insulin secretion (GSIS) of human β–cells Insulin granules in monolayer EndoC–βH1 cells

To verify functionality of human β–cell line, parental EndoC–βH1 cells was grown in two–dimensional monolayer by immunofluorescence microscopy (Figure 4). Cells were fixed 72 h after seeding, permeabilized, and stained for insulin, with nuclei counter–labelled using DAPI. Confocal imaging revealed intense insulin signal concentrated in the perinuclear cytoplasm of every cell, consistent with dense–core secretory granules typical of mature β–cells. The presence of discrete green granules confirms preserved pro–insulin processing and granule biogenesis which is prerequisite for physiologically relevant glucose–stimulated insulin secretion and validates EndoC–βH1 as an appropriate human model for downstream gene–editing and transplantation studies.

**Figure 4.**
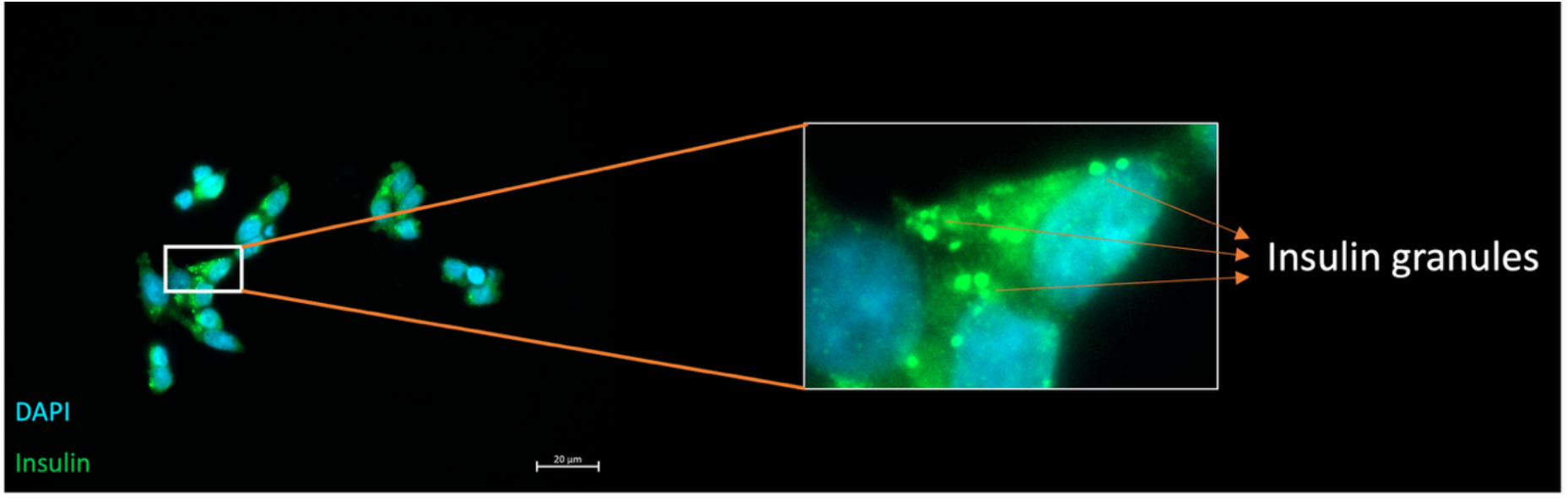
Immunofluorescence demonstration of insulin–containing granules in EndoC–βH1 cells. Representative confocal micrograph of unedited EndoC–βH1 cells cultured as a monolayer. Insulin immunoreactivity (green) appears as concentrated in the perinuclear region, while nuclei are counter–stained with DAPI (blue). The magnified inset highlights individual dense–core insulin granules (arrowheads), confirming intact endocrine granule biogenesis in the human β–cell line. (Scale bar = 20 µm)

### Insulin production in EndoC–βH1 spheroids

To visualize insulin production within the spheroids, immunofluorescent labeling was conducted. Hand–picked spheroids were centrifuged at 100 g for 5 minutes, and the pellet was fixed with 4% PFA for 15 minutes. Following washing and incubation with an insulin primary antibody, staining with DAPI for nuclei and Alexa Fluor 488 for insulin was performed. Visualization was carried out using an 8–well chambered glass slide under a confocal microscope (Figure 5). Each cell within the spheroids exhibited insulin production, further demonstrating the suitability of EndoC-βH1 spheroids for subsequent experiments.

**Figure 5.**
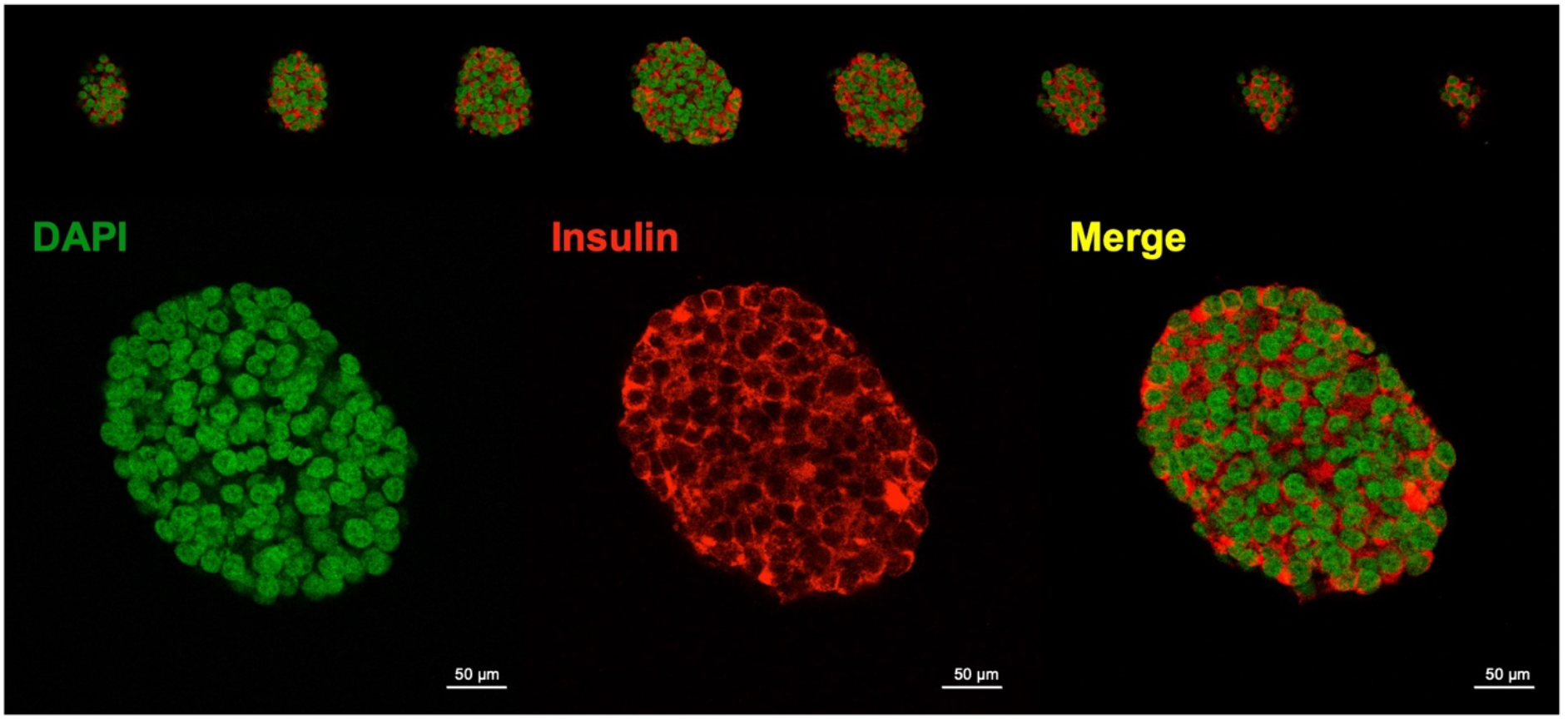
Confocal immunofluorescence of EndoC–βH1 spheroids confirms homogeneous insulin expression throughout the aggregate. Z–stack projections (top) and a single optical section (bottom) of 3–day–old EndoC–βH1 spheroids generated on agarose ultra–low–attachment dishes. Nuclei are stained with DAPI (green pseudocolour), insulin–positive dense–core granules (red), and the composite merge (yellow), indicating that essentially every cell within the aggregate retains a β–cell phenotype. (Scale bars = 50 µm)

### Glucose stimulated insulin secretion (GSIS) of B–TC–6 and EndoC–βH1 spheroids

Glucose-stimulated insulin secretion (GSIS) was next examined to determine whether gene editing affected β–cell function (Figure 6). In mouse β-TC-6 cells, the NT control exhibited a stimulation index (SI) close to 1, indicating that these cells are not responsive to different glucose challenges. Disruption of HIVEP2 had no impact, yielding the SI statistically non-significant from control. Additionally, the knockout of Ins2 resulted in the sharp decrease of insulin production. However, SI was calculated as 1 because the ratio of insulin at high glucose over low glucose is similar due to the disrupted insulin production mechanism (excluding unresponsiveness of murine β–cell). Notably, Rnls-edited cells displayed the SI fell by roughly 50 % relative to control, indicating that loss of Rnls compromises, but does not completely abolish, glucose-stimulated insulin release in the murine line (Figure 6a).

Human EndoC-βH1 cells showed the much higher SI characteristic of this more mature model. Again, HIVEP2 deletion left glucose responsiveness intact, whereas INS knockout dramatically suppressed secretion. In contrast to the mouse data, RNLS editing produced only a modest, statistically non-significant reduction in SI, suggesting that loss of RNLS was not compromises insulin secretion of human β–cells (Figure 6b). Together, these functional assays reinforce the idea that HIVEP2 deletion is not dampen insulin secretion, while Rnls loss exerts a cell-line-dependent decrease in insulin secretion, prominent in mouse β-TC-6 cells.

**Figure 6.**
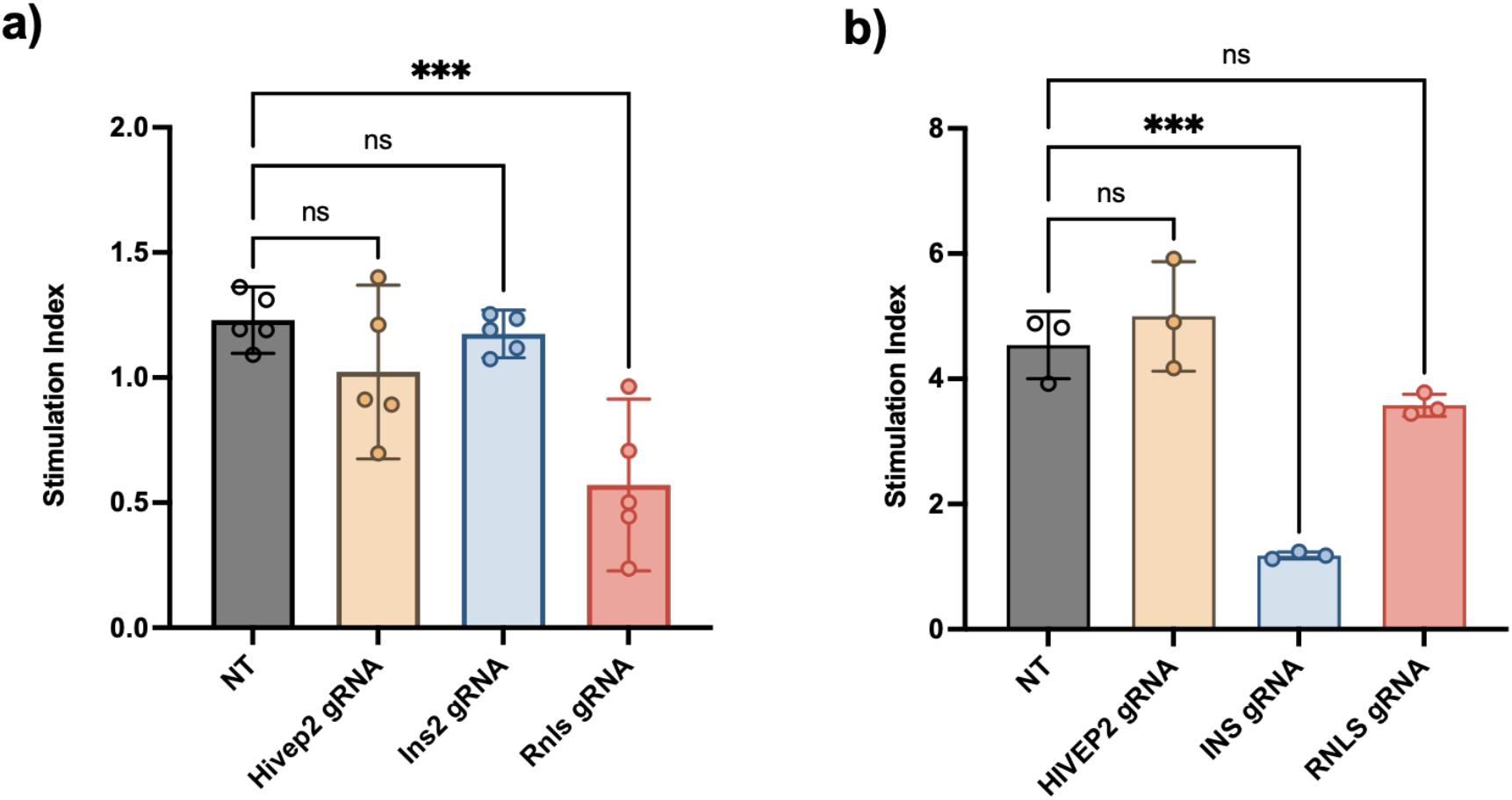
Glucose–stimulated insulin–secretion (GSIS) profiles of gene–edited β–cell spheroids. Stimulation index (ratio of insulin released at HG versus LG; mean ± SD, 30 spheroids for batches) for **a)** murine β–TC–6 and **b)** human EndoC–βH1 spheroids carrying non–targeting (NT) or gene–specific guides. In mouse cells, cells showed no response against different glucose challenges, whereas Hivep2 deletion was neutral and Rnls deletion produced a moderate but significant reduction (P < 0.05) relative to NT. In human cells, INS knockout again eliminated GSIS (*** P < 0.001) while neither HIVEP2 nor RNLS knockout altered the stimulation index (ns, not significant).

### In vivo transplantation

Before initiating the gene-knockout *in vivo* transplantation experiments, a pilot experiment was done to verify whether EndoC–βH1 spheroids can survive long-term in an immunodeficient environment. Luciferase-labelled spheroids (5 × 10^6^ cells, Matrigel embedded) were implanted subcutaneously into the right flank of either CD-1 mice (fully immunocompetent) or severe combined immunodeficiency disease (scid) mice (T and B cell deficient). Longitudinal bioluminescence imaging (d-luciferin, 150 mg/kg) revealed rapid loss of graft signal in CD-1 animals, falling below background by day 7, whereas identical grafts in scid mice persisted and expanded over a 35-day observation period (Figure 7). These data confirm that innate immune mechanisms alone are insufficient to eliminate the graft, and that adaptive immunity is the dominant driver of xenogeneic rejection in our system.

**Figure 7.**
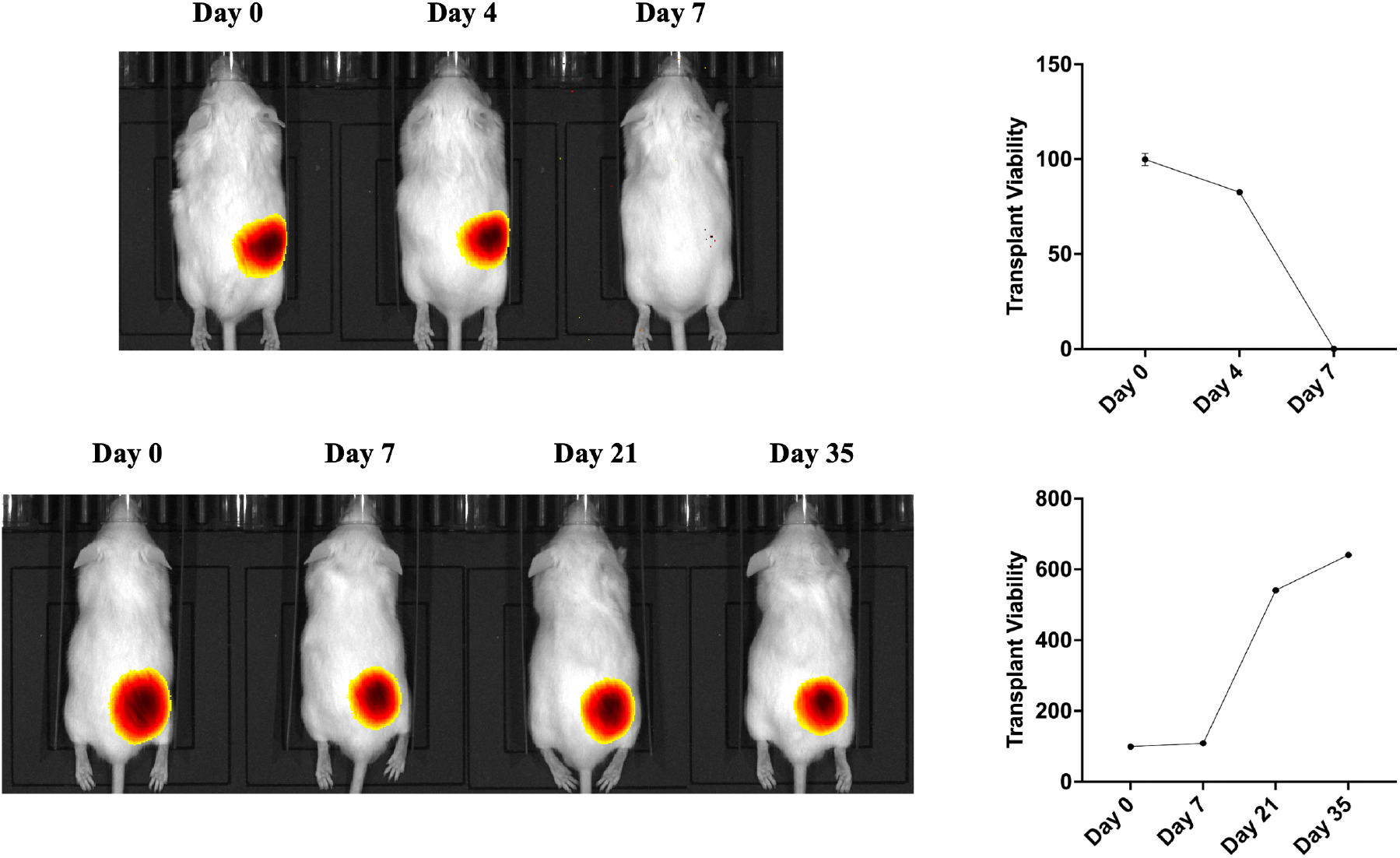
Human EndoC–βH1 graft fate in immunocompetent versus immunodeficient mice. Bioluminescence images of a CD-1 mouse (top) and a scid mouse (bottom) bearing identical luciferase-expressing EndoC–βH1 spheroids. Corresponding photon-flux curves (right) plot residual graft viability as a percentage of day-0 signals. In CD-1 mice the xenograft is rapidly rejected, with complete loss of luminescence by day 7. In contrast, the same graft in scid mice persists and progressively enlarges, demonstrating that adaptive immunity is required for xenogeneic clearance in this model.

### Allotransplantation of B–TC–6 spheroids to CD1 mice

To evaluate whether targeted gene knockouts influence β–cell survival in an allogeneic transplantation context, luciferase–labelled β–TC–6 cells genetically edited for Hivep2, Ins2, and Rnls, as well as NT controls, were transplanted subcutaneously into the flank region of immunocompetent CD1 mice using Matrigel as a matrix (Figure 8). Cellular survival was assessed through bioluminescence imaging (BLI) following intraperitoneal administration of D–luciferin. The Hivep2 knockout cells showed a similar survival trajectory compared to NT controls, with rapid signal loss evident by day 7, indicating no protective benefit against allogeneic immune rejection. Interestingly, Rnls knockout cells exhibited prolonged survival relative to NT controls, maintaining comparable bioluminescence intensity through day 10 and exhibiting a slower decline thereafter. Although these cells were eventually cleared, the delay suggests that RNLS deficiency partially improves beta–cell survival in the face of allogeneic rejection. These findings collectively reveal that both Rnls and Hivep2 did not affect beta–cell survival in allogeneic transplantation, indicating that Rnls protective effect on autoimmune attack does not translate in allorejection.

**Figure 8.**
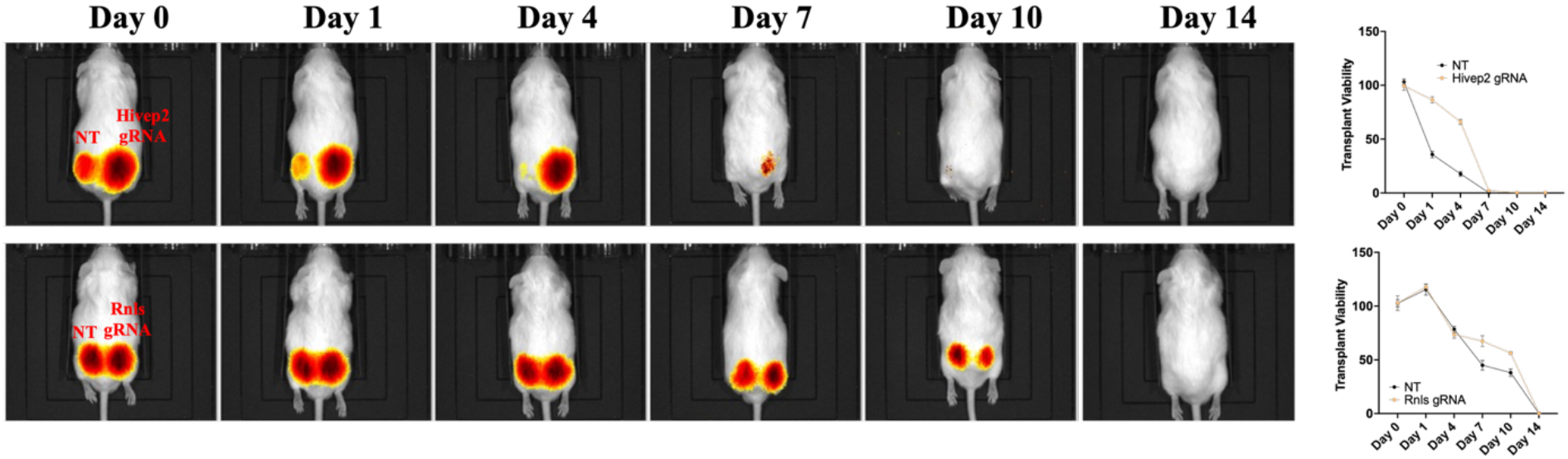
*In vivo* bioluminescence tracking of allogeneic β–TC–6 spheroid grafts in CD–1 mouse. Luciferase–expressing spheroids (5 × 10^6^ cell per flank, embedded in Matrigel) were implanted subcutaneously on day 0; left flanks carried a non–targeting (NT) control, right flanks received a gene–edited counterpart. Serial ventral images (days 0–14; identical colour scale) are shown for HIVEP2 (top) and Rnls (bottom) knockouts. Line plots on the right depict residual photon flux, expressed as percentage of day–0 signal, for NT (black) versus each edit (orange). All grafts declined progressively and were undetectable by day 14; neither HIVEP2 nor RNLS deletion altered the clearance curve.

### Xenotransplantation of EndoC–βH1 spheroids to CD1 mice

To test the effect of gene knockouts in xenotransplantation, luciferase-labelled EndoC–βH1 cells, harboring HIVEP2 or RNLS knockouts or a NT control, were embedded in Matrigel and implanted into flank region of immunocompetent CD-1 mice (Figure 9). Across all groups, signal decay was markedly faster than in the allogeneic Beta-TC-6 transplantation, reflecting the increased severity of xenorejection. HIVEP2-edited grafts lost >80 % of their luminescence by day 4 and were undetectable by day 7, indicating no xenoprotective effect.

**Figure 9.**
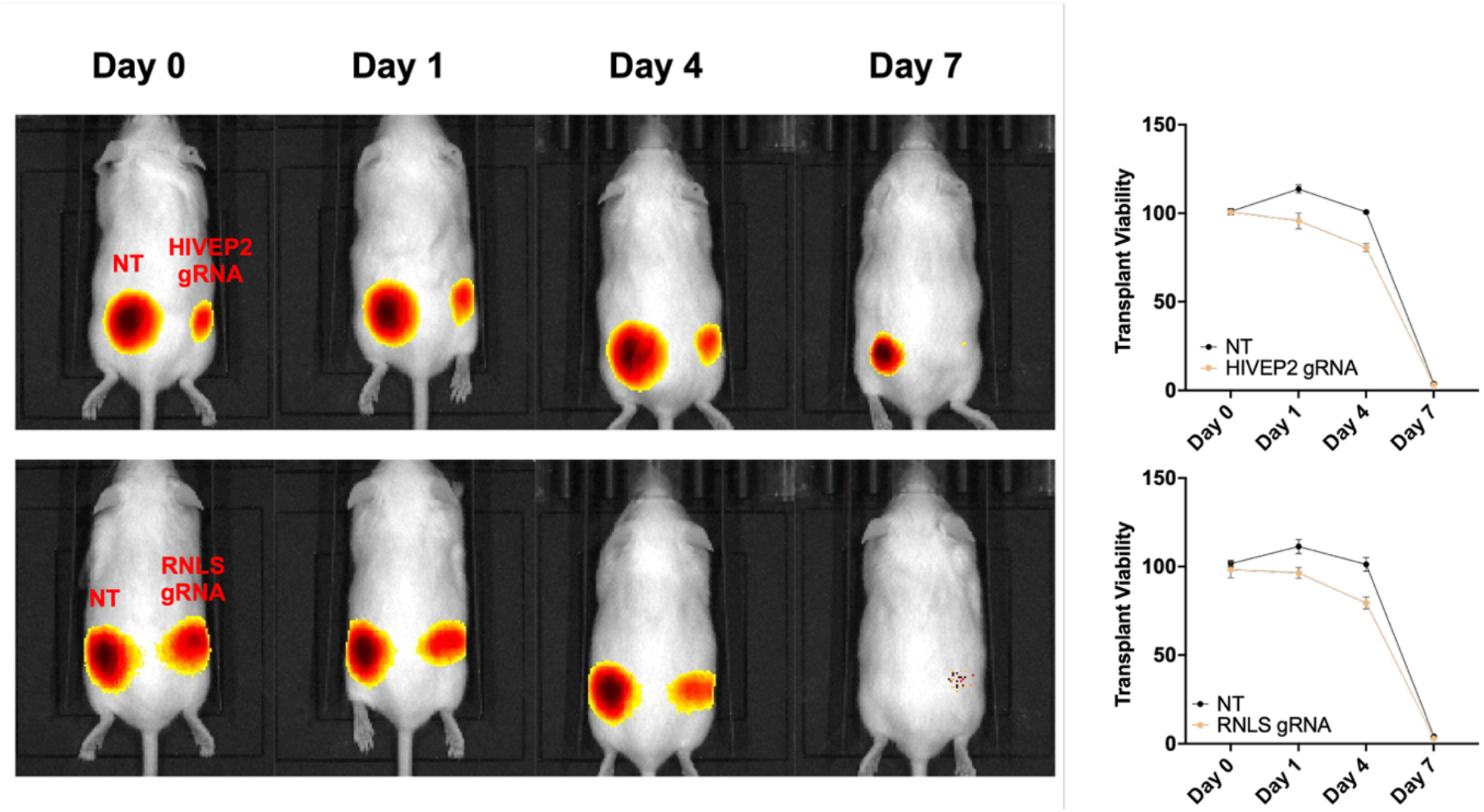
Rapid xenogeneic rejection of EndoC–βH1 spheroid grafts in CD–1 mouse. Luciferase–labelled human EndoC–βH1 spheroids (5 × 10^6^ cells per flank, Matrigel embedded) were implanted subcutaneously on day 0; the left flank received a non–targeting (NT) control, the right flank the indicated CRISPR knockout. Sequential bioluminescence images are shown for HIVEP2 (top) and RNLS (bottom) edits, together with quantitative photon–flux decay curves (right; percentage of day–0 signal). All grafts, irrespective of genotype, lost >80 % of baseline radiance by day 4 and were completely eliminated by day 7, illustrating the formidable innate and adaptive barriers to human–to–mouse xenotransplantation. Neither RNLS nor HIVEP2 knockout conferred any detectable survival advantage and did not alter rejection kinetics.

RNLS-edited grafts displayed a modest delay in clearance, retaining ~30 % of baseline signal on day 7 versus <10 % in controls, but were nonetheless fully rejected by day 10. Thus, in the xenogeneic CD-1 model, none of the single-gene edits conferred durable protection to human β–cells. These findings underline the greater immunological hurdle posed by xenotransplantation and suggest that more extensive engineering, potentially combining RNLS or HIVEP2 loss with additional immune-evasion strategies, will be required to achieve long-term engraftment of human β–cells in fully immunocompetent hosts.

### Conclusion and future directions

This work provides the first direct evaluation of RNLS and HIVEP2 knock–out as stand–alone immune–evasion strategies outside an autoimmune context. By combining high–viability, size–controlled β–cell spheroids with rigorous genomic and functional validation, we show that neither edit prolongs graft survival in fully immunocompetent recipients whether across an allogeneic (mouse → mouse) or xenogeneic (human → mouse) barrier. RNLS deletion shows a modest insulin–secretory reduction in murine cells, whereas HIVEP2 loss is functionally neutral, yet neither modification alters the inevitable immune clearance observed *in vivo*. These findings set an upper bound on the protective capacity of these loci when targeted in isolation and emphasize the complexity of transplant immunity relative to recurrent autoimmunity.

The results obtained in this study highlight several critical insights into the complexity of immune rejection processes [35, 36]. One potential reason for the failure could be that immune rejection involves redundant and compensatory mechanisms, such that disrupting only a single regulatory gene is insufficient to halt the complex interplay of immune responses. Additionally, while RNLS or HIVEP2 regulate apoptosis and immune signaling pathways, other dominant pathways or immune effector mechanisms may still mediate effective graft destruction, dominating any minor protective effect provided by the single knockout. These results emphasize the necessity for multi-gene editing approaches, suggesting that future genetic modifications should target multiple pathways simultaneously such as combining disruption of antigen presentation (HLA molecules) with expression of immunomodulatory molecules (PD-L1, CD47). Moreover, findings stress the importance of selecting genetic targets based not only on their isolated molecular roles but also considering their broader network interactions within the cellular and immune microenvironment.

The study nevertheless establishes several assets for future engineering efforts: (i) a scalable agarose suspension platform that yields uniformly sized, viable spheroids well suited to transplantation and imaging; (ii) a CRISPR workflow achieving higher editing efficiency in both mouse and human β–cells; and (iii) a quantitative bioluminescence pipeline that permits head–to–head comparison of candidate edits across allo– and xeno–models.

Moving forward, durable engraftment will almost certainly require combinatorial approaches [24, 37]. Rational strategies include HLA class I/II elimination, “don’t–eat–me” signals (CD47, PD–L1), or inducible suicide switches to minimize oncogenic risk [38]. Parallel advances in biomaterials including macro– or micro-encapsulation [39, 40], 3-D bioprinting [41, 42], and local immune–modulating hydrogels [43] could synergize with multi–gene editing approach. Finally, humanized–immune and complement–sufficient models may provide a more predictive test bed for such next–generation β–cell products. By defining the limitations of single–gene edits, our study supplies a critical benchmark and an experimental framework for the layered engineering that will be required to translate β–cell replacement into a practical, immunosuppression–free cure for T1D.

## Materials and Methods

### Cell lines

β–TC–6 and 293T (CRL–3605 and CRL–3216) cell lines were obtained from ATCC, EndoC–βH1 cell line was obtained from Human Cell Designer. 293T cells were maintained in DMEM (D6429, Sigma), supplemented with 10% FBS (S1520, Biowest), and penicillin/streptomycin (L0022, Biowest) in a 37 °C incubator with 5% CO2. β–TC–6 cells were maintained in low bicarbonate DMEM (P04–03596, PAN Biotech), supplemented with 15% FBS (S1520, Biowest), and penicillin/streptomycin (L0022, Biowest) in a 37 °C incubator with 5% CO2. EndoC–βH1 cells were maintained in Opti–β1 (Human Cell Design) after coating with βcoat (Human Cell Design) in a 37 °C incubator with 5% CO2.

### gRNA Cloning

To generate NT, Hivep2KO, Ins2KO, and RnlsKO β–TC–6 cells, three gRNAs were selected from Brie mouse gRNA database (Supplementary Table 1) [44] and cloned into pLentiCRISPRv2 vector. After an initial screen of T7EI and TIDE analysis, one gRNA was selected for each gene as follows: NT gRNA (5′–GCGCTTCCGCGGCCCG–3′), Hivep2 gRNA (5′–TAAGGCGGATGACTCTCACA–3′), Ins2 gRNA (5′–GGACTCCCAGAGGAAGAGCA–3′), and Rnls gRNA (5′–CTACTCCTCTCGCTATGCTC–3′) were cloned into pLentiCRISPRv2 vector with puromycin antibiotic resistance gene (Addgene #98290) [45]. The gRNA containing lentiviruses were used to establish these cell lines.

To generate NT, HIVEP2KO, INSKO, and RNLSKO EndoC–βH1 cells, similarly, three gRNAs were selected from Brunello human gRNA database (Supplementary Table 5) [44] and cloned into pLentiCRISPRv2 vector. After an initial screen of T7EI and TIDE analysis, one gRNA was selected for each gene as follows: NT gRNA (5′–GACGGAGGCTAAGCGT–3′), HIVEP2 gRNA (5′–GACAAGATGTCAGACCTAGG–3′), INS gRNA (5′–GAAGCTCTCTACCTAGTGTG–3′), and RNLS gRNA (5′–TCCCACACAGCAAGGTACAA–3′) were cloned into pLentiCRISPRv2 vector with puromycin antibiotic resistance gene. All plasmid sequences were verified by Sanger sequencing before transduction and transfection (Supplementary Figure 1).

### Lentivirus production

293T cells were seeded at 4.0 × 106 cells on a 10–cm plate 1 day before transfection. The cells were transfected with 2.5 μg of a vector, 2.25 μg of a transfer plasmid (psPAX2, #12260; Addgene), and 0.25 μg of a VSV–G vector (pMD2.G, #12259; Addgene) using the polyethylene imine (PEI) transfection reagent (Thermo Fisher Scientific, Waltham, MA). Viral particles were harvested 48 h post transfection and purified using a 0.45–μm Millex–HV filter (SLZHVR33RS; Merck Millipore, Burlington, MA).

### Virus infection and drug selection

For viral transduction, 5 × 10^6^ cells were seeded in 10–cm tissue–culture dishes and allowed to attach for 24 h. The cultures were then exposed overnight to 10 µL of 100–fold–concentrated lentiviral stock supplemented with 10 µg/mL protamine sulfate (P3369, Sigma–Aldrich) to enhance viral adsorption. Fresh growth medium was provided the next morning. Forty–eight hours later, robust fluorescence in parallel GFP–virus controls confirmed successful transduction, and antibiotic selection was initiated. β–TC–6 cultures received 4.5 µg/mL puromycin, whereas EndoC–βH1 cultures were treated with 2.5 µg/mL. Selection was maintained for four days, after which surviving cells were expanded for an additional two days in antibiotic–free medium to recover. All downstream assays were performed with these fully selected, puromycin–resistant populations.

### T7 endonuclease I (T7E1) assay

CRISPR–Cas9 editing in both β–TC–6 and EndoC–βH1 cells were detected by T7 endonuclease I mismatch cleavage assay. Genomic DNA was purified from NT and gRNA transduced cells using the Macherey–Nagel™ NucleoSpin™ Tissue, Mini kit for DNA from cells and tissue. The Hivep2, Rnls, and insulin gRNA targeting site was amplified using the Phusion High–Fidelity PCR Kit. The primers for the gRNA site PCR were as follows: Hivep2 site forward, 5′–CTCCCTGCTGAGAAGTTGCC–3′; reverse, 5′–GATGTGGCTGTTCGGGTAAG –3′, Ins2 site forward, 5′–CGTGAAGTGGAGGACCCACA–3′; reverse, 5′–AAACTGTGGGTCCTCCACTTCACG–3′, Rnls site forward, 5′–TATCCCATGTGGCTTGGAGT–3′; reverse, 5′–GAGGTGAAGAATCGGTCCACT–3′, HIVEP2 site forward, 5′–GTTGTGTTGCCGAACTAGCC–3′; reverse, 5′–GGGGACAGGGTTGGGTATGA–3′, INS site forward, 5′–CCACCCTCTGATGTATCTCGG–3′; reverse, 5′–AGACTATAAAGCCAGCGGGG–3′, RNLS site forward, 5′–GAGTGGAATCAATCTTAGCAGTTG–3′; reverse, 5′–GAGTGGAATCAATCTTAGCAGTTG–3′. The 400 ng PCR products were used to form heteroduplexes by denaturing at 95 °C for 5 min and then reannealing the products in a thermocycler using the following protocol: ramp down to 85 °C at −2 °C per s; ramp down to 25 °C at −0.1 °C per s; hold at 4 °C. Ten units of T7 endonuclease I (#M0302) were added to the annealed PCR products and the reaction was incubated at 37 °C for 15 min. The digestion reaction was stopped by 1 μl of 0.5 M EDTA and immediately applied to a 1% agarose gel to visualize digested and undigested products by electrophoresis.

### Hanging–drop aggregation

Single–cell suspensions were prepared in complete growth medium (DMEM, 10 % FBS, 1 % penicillin/streptomycin) at 1 × 10^7^ cells/mL. Thirty–microlitre droplets containing either 300, 500, or 1000 cells were dispensed onto the inner surface of 100–mm Petri–dish lids. Lids were inverted over dishes containing 10 mL sterile PBS to maintain humidity and incubated for 72 h at 37 °C, 5 % CO_2_. Spheroids that formed at the bottom of the droplet were harvested by gentle pipetting with pre–warmed medium.

### Home-made Ultra–low–attachment (ULA) suspension culture

For higher throughput, homemade ULA dishes were fabricated by pouring 5 mL of sterile 1 % (w/v) agarose in PBS into 100–mm Petri dishes, allowing the gel to solidify for 30 min, and rinsing once with growth medium. Subsequently, 5 × 10^6^ single cells in 10 mL medium were added per dish and cultured on an orbital shaker (50 rpm) for 3 days at 37 °C, 5 % CO_2_. Gentle agitation promoted rapid self–assembly into compact spheroids with minimal size heterogeneity.

### Live/Dead staining and imaging

Spheroids from both protocols were washed twice with warm PBS and incubated for 3 min with fluorescein diacetate (FDA, 2 µg/mL) at 37 °C to label metabolically active cells. Propidium iodide (PI, 1 µg/mL) was then added for an additional 5 min to stain membrane–compromised cells. After two PBS washes, spheroids were transferred to glass–bottom dishes and imaged immediately by epifluorescence microscopy (FITC and Texas Red filter sets). Aggregate diameters were measured from bright–field micrographs using ImageJ.

### Glucose stimulated insulin secretion (GSIS)

For glucose–stimulated insulin secretion assay, the cells were plated in a 6–well plate as 1.5 × 106 cells per well. After 48h, cells were washed twice very gently using Krebs–Ringer Buffer (KRB). Before starting GSIS, cells were starved for 60 minutes by leaving cells in 2 mL of KRB buffer and incubating at 37°C, 5% CO2. After starvation, KRB buffer was removed and cells were incubated using 2.8 mM D–Glucose KRB (low glucose, LG) at 37°C, 5% CO2 for 60 minutes. Supernatant was collected and then cells were finally incubated using 16.7 mM D–glucose KRB (high glucose, HG) for 60 minutes at 37°C, 5% CO2, and the supernatant was collected. Supernatant was used to perform insulin quantification using the insulin ELISA kit from Mercodia.

For β–TC–6 the KRB contained (in mmol/L): 115 NaCl, 5.3 KCl, 2.6 CaCl_2_, 1.2 MgSO_4_, 1.0 KH_2_PO_4_, 24 NaHCO_3_ and 10 HEPES, supplemented with 0.1 % (w/v) fatty–acid–free BSA and either 1.67 mM (LG) or 16.7 mM (HG) glucose. The solution was equilibrated in 5 % CO_2_ to maintain pH 7.35–7.40.

For EndoC–βH1 the KRB containing (in mmol/L): 138 NaCl, 5.4 KCl, 2.6 CaCl_2_, 0.5 MgCl_2_, 5 NaHCO_3_ and 10 HEPES, again with 0.1 % fatty–acid–free BSA and either 2.8 mM (LG) or 28 mM (HG) glucose.

### In vivo imaging (IVIS)

To enable *in vivo* bioluminescence tracking, both β–cell lines were first transduced with a lentiviral construct with the firefly luciferase (Fluc). Fluc coding sequence is driven by the human ubiquitin–C (ubC) promoter with a selection marker of Neomycin (G418) or hygromycin B (Supplementary Figure 14). Stable cell lines were obtained by antibiotic selection, 400 µg/mL G418 for β–TC–6 cells and 200 µg/mL hygromycin B for EndoC–βH1 cells, resulting in uniform Fluc expression.

For *in vivo* studies, 5 × 10^6^ Fluc–cells from each knockout line, together with a non–targeting (NT) control, were resuspended in 200 µL Matrigel (CLS354234, Merck) and injected subcutaneously into the left and right flank of CD–1 mice (designated Day 0). Bioluminescence imaging (BLI) was performed with a Caliper IVIS Spectrum system (PerkinElmer). Ten minutes before image acquisition, animals received an intraperitoneal injection of d–luciferin (150 mg/kg of animal; P1042, Promega) prepared at 15 mg/mL in calcium– and magnesium–free Dulbecco’s PBS and sterile–filtered through a 0.22–µm membrane. Imaging parameters were kept constant throughout the study (exposure = 1 min, binning = medium, f/stop = 1, field of view = 12.5 cm) to permit quantitative comparison. Photons emitted from the graft region were integrated with Living Image software and expressed as radiance (photons/scm^2^sr).

### Animal experiments

All *in vivo* procedures were approved by the Koç University Animal Research Facility Ethical Committee (KUARF, approval no. 2024.HADYEK.038) and conducted in accordance with the ARRIVE guidelines and Turkish Animal Welfare regulations. Female CD–1 or SCID mice (8–10 weeks, 25–30 g) were housed in individually ventilated cages (22 ± 2 °C, 50 ± 10 % humidity, 12 h light/dark cycle) with ad libitum access to standard chow and water.

For each genetic condition, NT control, Hivep2/HIVEP2 knockout, Rnls/RNLS knockout, and Ins2/INS knockout, n = 4 mice were used in both the allogeneic (β–TC–6 → CD–1) and xenogeneic (EndoC–βH1 → CD–1) cohorts. On day 0 mice were anaesthetised with 2 % isoflurane, and spheroids from 5 × 10^6^ luciferase–expressing cells suspended in 200 µL growth–factor–reduced Matrigel were injected subcutaneously into each flank (left, NT; right, gene–edited) and visualized via IVIS. Mice were euthanized by CO_2_ inhalation followed by cervical dislocation.

### Statistical analysis

One–way ANOVA with Tukey comparison test on GraphPad Prism Software (Graph–Pad Software version 10.5, CA, USA) was used. A value of p ≤ 0.05 was considered statistically significant. The significance levels are denoted as follows: n.s. for p > 0.05, * for p ≤ 0.05, ** for p ≤ 0.01, *** for p ≤ 0.001, and **** for p ≤ 0.0001.

## Acknowledgement

The authors sincerely acknowledge the use of the facilities and services provided by the Koç University Research Center for Translational Medicine (KUTTAM). The financial support for this project was provided by the Scientific and Technological Research Council of Turkey (TUBITAK) under an International Support Program (COST Action - European Cooperation in Science and Technology− CA20140, project number: 122S968). I.C.K. would like to acknowledge BIDEB scholarships from TUBITAK.

## Author contributions

I.C.K. and S.K. conceived the idea and S.K. supervised the project together with T.O.. I.C.K. performed all experiments with support from A.O. (for cloning experiments). The manuscript was written by I.C.K. and S.K.. All authors read and approved the final version of the manuscript.

## Notes

### Competing Interest Statement

The authors have declared no competing interest.

