## Supplementary material for "Single–gene knockout of RNLS or HIVEP2 are insufficient to protect β–cell spheroids from allo– and xeno–rejection": supp-info

**Supplementary Table 1. gRNA sequences for targeting Hivep2, Ins, and Rnls genes in both mouse and human insulin secreting beta cells.**

The overhanging sequences highlighted with green for integration to pLentiCRISPRv2 plasmid after BsmBI digestion.

| | | 5' – 3' (Top oligo) | 3' – 5' (Bottom oligo) |
|---|---|---|---|
| **Mouse** | Hivep2_1 | CACCGCCTGTGAGAACAG AAAACGC | CGGACACTCTTGTCTTTT GCGCAAA |
| | Hivep2_2 | CACCGTAAGGCGGATGAC TCTCACA | CATTCCGCCTACTGAGAG TGTCAAA |
| | Hivep2_3 | CACCGACAATGAGTATGA ACCCGCA | CTGTTACTCATACTTGGGC GTCAAA |
| | Ins2_1 | CACCGGTGGAACAACTGG AGCTGGG | CCACCTTGTTGACCTCGA CCCCAAA |
| | Ins2_2 | CACCGTAGAGAGCCTCTA CCAGGTG | CATCTCTCGGAGATGGTC CACCAAA |
| | Ins2_3 | CACCGCTGGGAGCCCAAA CCCACCC | CGACCCTCGGGTTTGGGT GGGCAAA |
| | Rnls_1 | CACCGCAGTAATTAGTGA ACGCCAG | CGTCATTAATCACTTGCG GTCCAAA |
| | Rnls_2 | CACCGCCACCTGAAAGAT AACAAGT | CGGTGGACTTTCTATTGTT CACAAA |
| | Rnls_3* | CACCGCTACTCCTCTCGC TATGCTC | CGATGAGGAGAGCGATAC GAGCAAA |
| **Human** | HIVEP2_1 | CACCGGACAAGATGTCAG ACCTAGG | CCTGTTCTACAGTCTGGA TCCCAAA |
| | HIVEP2_2 | CACCGTTCTAGGATAACC ACCACTG | CAAGATCCTATTGGTGGT GACCAAA |
| | HIVEP2_3 | CACCGGAGGTGGAAGGT AAACACAA | CCTCCACCTTCCATTTGTG TTCAAA |
| | INS_1 | CACCGGCACAGGTGTTGG TTCACAA | CCGTGTCCACAACCAAGT GTTCAAA |
| | INS_2 | CACCGCGGGAGGCAGAG GACCTGCA | CGCCCTCCGTCTCCTGGA CGTCAAA |
| | INS_3 | CACCGGAAGCTCTCTACC TAGTGTG | CCTTCGAGAGATGGATCA CACCAAA |
| | RNLS_1 | CACCGATTATGCCAAAAA ACACCAA | CTAATACGGTTTTTTGTGG TTCAAA |
| | RNLS_2 | CACCGTCCCACACAGCAA GGTACAA | CAGGGTGTGTCGTTCCAT GTTCAAA |
| | RNLS_3 | CACCGCATTACTTGAAAG AATCAGG | CGTAATGAACTTTCTTAGT CCCAAA |

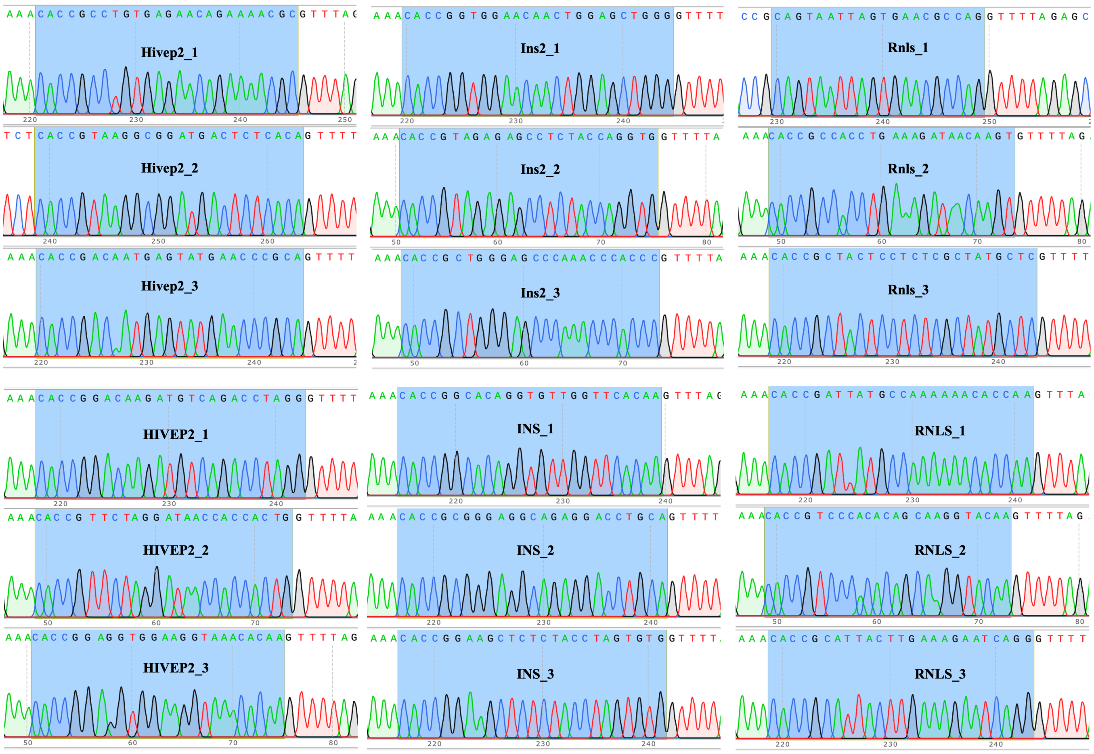


**Supplementary Figure 1. Sanger sequencing confirmation of individual gRNA inserts in the pLentiCRISPRv2 vector backbone.**

Sanger sequencing validations for gRNA cloning into lentiCRISPRv2 plasmid. (Hivep2, Ins2, Rnls for mouse cells while HIVEP2, INS, RNLS for human cells.)
